# Inter- and intra-specific variation in the pelagic larval duration of four Tropical Eastern Pacific damselfishes (Pomacentridae: *Stegastes*) with contrasting distribution patterns

**DOI:** 10.1101/2024.08.31.610633

**Authors:** D. F. Lozano-Cortés, M. Rodríguez-Moreno, F. A. Zapata

## Abstract

Dispersal is a key ecological function in marine populations that are naturally fragmented and sometimes isolated. Pelagic larval duration (PLD) is thought to approximate the dispersal potential of coral reef fishes and may reflect the extent of connectivity in reef fish populations. Inter- and intra-specific variation in PLD of four damselfish species was investigated in the Colombian Pacific at four locations (two on the mainland coast, one at a continental island and one at an oceanic island) at multiple spatial scales (between localities and between sites within a single locality). Two of the species (*Stegastes acapulcoensis* and *S. flavilatus*) are broadly distributed on the continental coast while the other two (*S. beebei* and *S. arcifrons*) are largely restricted to oceanic islands. Nonetheless, individuals of continental species sporadically colonize oceanic habitats, and vice-versa. The PLD was estimated using counts of otolith growth increments from juveniles collected at all localities. Species with an oceanic distribution had longer PLDs than their congeners with continental distributions. Differences in PLD between the two continental species varied between localities and significant intra- specific spatial variability was observed between localities but not within a single locality. Although the species studied have the necessary PLD to reach all available habitat, there are no apparent colonization events between mainland and oceanic islands suggesting that their distribution is not limited by dispersal but by other processes. We discuss the possible underlying causes of the observed variability, and suggest the need to consider spatial variability in the development of dispersal models and connectivity patterns for better management of coral reef fish populations.

## INTRODUCTION

The life cycle of most reef fishes includes a pelagic larval phase and a benthic adult phase (Leis, 1991). The larval phase begins when the larva hatches from the egg, and ends days or weeks later when the larva undergoes metamorphosis and settles on the reef. Because the benthic adults are relatively sedentary and reefs are patchily distributed, dispersal and colonization of new habitats occurs primarily during the pelagic larval phase (Leis, 1991; Shanks, 2009; Cowen & Sponaugle, 2009). While it is likely that some dispersal is achieved through larval transport by currents (White *et al*., 2010), larvae also have strong swimming, behavioral and orientation abilities that may influence their dispersal (Leis *et al*., 1996; Leis & McCormick, 2002; Paris & Cowen, 2004; Fisher, 2005). The time that the larvae spend in the plankton may determine the dispersal potential of reef fishes, thus the longer the pelagic larval duration (hereafter PLD), the greater the possibilities of dispersing farther and colonizing remote habitats (Shanks *et al*., 2003).

In marine fishes, PLD can be estimated through the analysis of the daily growth increments formed in the otoliths of juveniles (Panella, 1971; Brothers *et al*., 1976). This information has shown to be consistent and reliable (Victor, 1982; Campana, 2001) and commonly used to study early life history processes such as larval dispersal, recruitment dynamics, early growth and survival of larvae. This knowledge therefore has important implications for understanding diverse ecological, biogeographical and evolutionary processes (Sponaugle, 2009).

Considerable variability in PLD at different taxonomic, temporal and spatial scales has been found in reef fishes (Victor, 1986; Thresher & Brothers, 1989; Wellington & Victor, 1989; Wellington & Victor, 1992; McCormick, 1994; Cowen & Sponaugle, 1997; Lester *et al*., 2007; Leis *et al*., 2013). Studies on inter-specific differences in PLD have been driven by interest in testing the hypothesis that geographic distribution is largely determined by dispersal capability. Support for this hypothesis, however, has generally been weak (Brothers & Thresher, 1985; Victor 1986; Thresher & Brothers, 1989; Thresher *et al*., 1989; Wellington & Victor, 1989; Victor & Wellington, 2000; Zapata & Herrón, 2002; Lessios & Robertson, 2006; Lester *et al*., 2007; Mora *et al*., 2012; Luiz *et al*., 2013). At the intra-specific level, PLD can vary significantly depending on the location or geographical region being examined (Victor, 1986; Thresher *et al*., 1989; Wellington and Victor, 1989; Thorrold & Milicich, 1990; Wellington & Victor, 1992; McCormick, 1994; Bay *et al*., 2006). Variation at this level has revealed spatial patterns in larval characteristics, such as growth and dispersal (e.g., Victor & Wellington, 1992), that provide a valuable context to understand processes that vary at geographical scales and that affect the life history of the pelagic larvae (Cowen & Sponaugle, 1997; Searcy & Sponaugle, 2000).

Examining spatial variation is also relevant in studies that intend to use PLD estimates to predict genetic differences among populations and develop meta- population models (Shanks, 2009), even though there is debate over whether or not PLD and genetic metrics reflect scales of dispersal (Weersing & Toonen, 2009; Selkoe & Toonen, 2011; Faurby & Barber, 2012; Dawson, 2014). However, most PLD studies are spatially and temporally limited because their estimates are based on data obtained during short periods of time from one or few locations. Increased data can lead to better estimates of important parameters and provide a more robust view of particular processes, therefore the need to study intraspecific variation in PLD, especially temporal and spatial variation, has gained importance. Recent studies have sought to fill this gap, both in temperate (Di Franco & Guidetti, 2011; Di Franco *et al*., 2013) and tropical regions (Bay *et al*., 2006; Booth & Parkinson, 2011; Kingsford *et al*., 2011).

In the Tropical Eastern Pacific (TEP) region, reef environments are patchily distributed along the continental coast and five oceanic islands or archipelagos, which include some of the smallest and most remote islands in the tropics. In their pioneering work on intraspecific variation in PLD, Wellington & Victor (1992) examined regional variation in the PLD of three species of reef fishes whose populations were geographically separated by 800 to 3500 km, and included populations from oceanic islands and continental localities. They found large regional differences in PLD, including differences between oceanic and continental populations.

In this study we examine inter- and intra-specific variation in PLD within and among populations of four congeneric species of damselfishes (Pomacentridae: *Stegastes*). Data was obtained from one oceanic island and three continental locations in the Colombian Pacific Ocean, separated by distances between 140 and 520 km, which represents a spatial scale where we might expect to find variability in PLD. The species considered exhibit contrasting patterns of geographic distribution: *Stegastes beebei* (Nichols 1924) and *S. arcifrons* (Heller & Snodgrass 1903) are largely restricted to oceanic islands (Galapagos Archipelago, Cocos Island and Malpelo Island), *S. acapulcoensis* (Fowler 1944) and *S. flavilatus* (Gill 1862) are broadly distributed on the continental coast of America from Mexico to Perú, and are absent or rare on oceanic islands (Robertson & Allen, 2015).

Nonetheless, some individuals of the continental species sporadically reach oceanic reefs, and vice-versa (Grove & Lavenberg, 1997; Victor & Wellington, 2000; Rodríguez-Moreno *et al*., 2011; Robertson & Allen, 2015) providing a rare opportunity to compare PLD among these fish species in relation to inferred dispersal events. Based on published studies related to fish PLD, dispersal and geographic distribution (e.g. tropical reef fishes in the Indo-Pacific; Lester & Ruttenberg, 2005), we expected longer PLDs in the species with larger distributions. Specifically, we asked 1) what are the differences in PLD among species? 2) Are inter-specific differences in PLD consistently related to oceanic *vs.* continental geographic distributions? More precisely, do similarly distributed species pairs exhibit similar PLDs? Furthermore, are inter-specific differences independent of location? 3) What is the extent of intra-specific spatial variation in PLD and how much of that variation is due to differences a) between oceanic *vs* continental localities, b) among localities within continental settings and c) among sites within a single locality?

## MATERIAL AND METHODS

### Study areas

This study considered four different localities (Fig. 1); two on the mainland, Málaga and Utría, separated by 247 km, and two on islands, Gorgona (a continental island 30 km off the mainland) and Malpelo (an oceanic island 380 km off the Colombian mainland), separated from each other by 396 km. Localities were visited one to three times between 2010 and 2013. Juveniles of the oceanic species (*S. beebei* and *S. arcifrons*) were observed and collected only at Malpelo Island and were not found at continental localities (although adults of *S. arcifrons* are not rare at Gorgona Island). In contrast, juveniles of the continental species *S. acapulcoensis* and S. *flavilatus* were collected at Malpelo Island. Although only one juvenile of *S. flavilatus* was collected at this locality, several juveniles of *S. acapulcoensis* were observed and collected at Malpelo Island.

**Figure 1.**
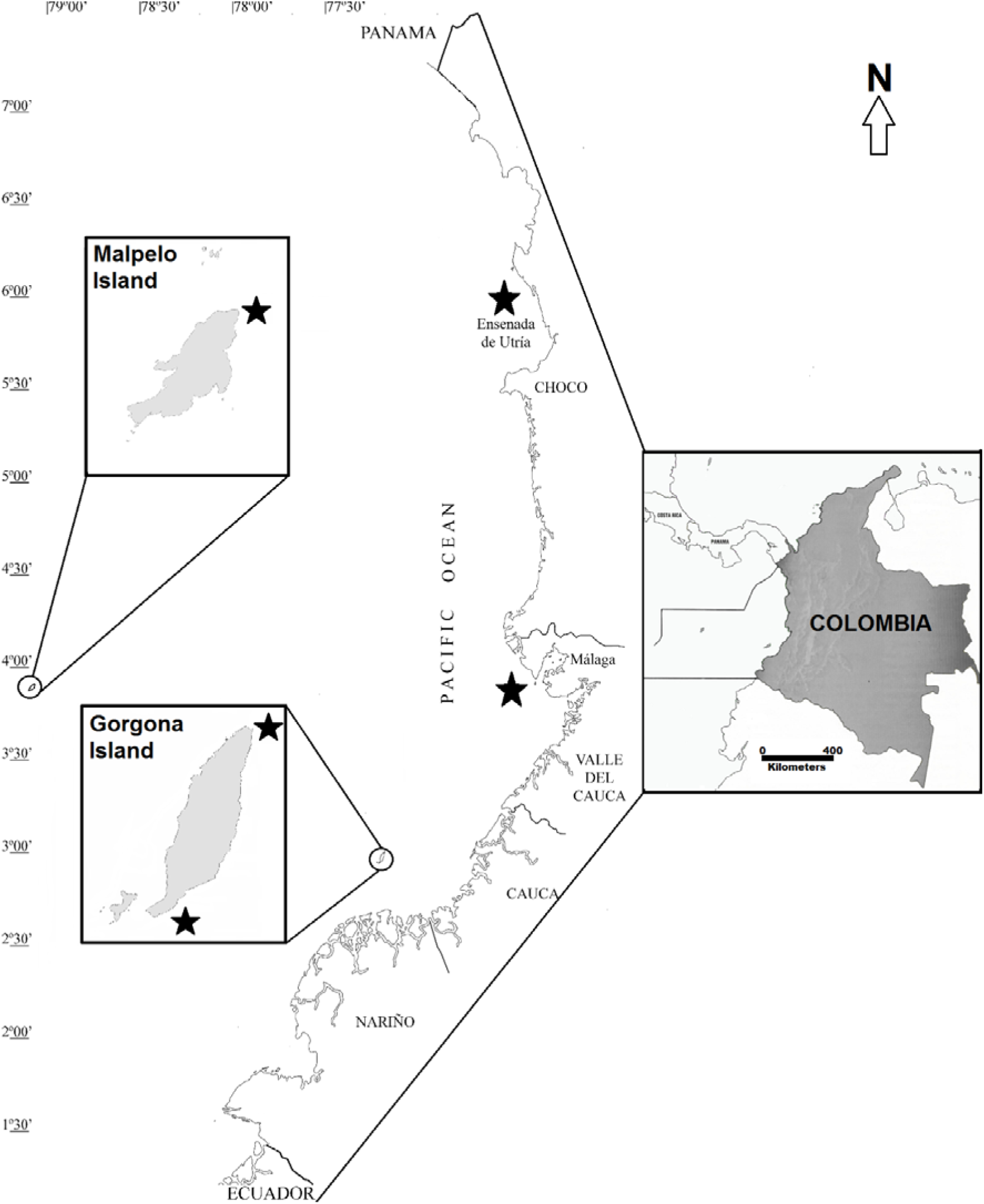
Map of the Colombian Pacific Ocean showing sampling locations.

Collections at Gorgona Island were carried out at two sites separated by 8.5 km (north and south of the island; Fig. 1). La Ventana is a coral community dominated by dense stands of *Pocillopora* spp. located in the south-eastern coast of Gorgona, while El Laberinto is a rocky reef located in the northernmost tip of the island. More detailed information from each locality is given by Zapata & Vargas-Angel (2003). During the first sampling period (November 2010 – February 2011), oceanographic conditions in the region corresponded to a cold episode of the Oceanic Niño Index (ONI), whereas during the second sampling period (July 2012 – February 2013) a neutral/weak ONI condition was reported (NOAA 2013).

Fish collection and otolith processing

Juvenile damselfishes were captured between 5 and 25 m depth in coral or rocky habitats at each location using a mixture of 70% ethanol and clove oil (3:1 ratio; Ackerman & Bellwood, 2002) and hand nets. A total of 190 juvenile fishes were collected (11.0 - 31.1 mm Standard Length), tagged and preserved in 95% ethanol for processing. Otolith extraction was carried out according to Secor *et al*. (1991, 1992) and Sponaugle (2009) under a stereomicroscope using tweezers via the operculum or through a head dissection. Subsequently, sagittae were fixed on the edge of a slide with thermoplastic glue (Crystal Bond). To get a transverse section, otoliths were ground with fine (1000-1200 grit) sandpaper from both distal and rostral ends and polished with 5.0 μm and 0.02 μm lapping film (3M).

PLD was estimated through the analysis of daily increments deposited in the otoliths during the pre-settlement stage (Sponaugle, 2009). The same reader counted the number of increments present from the primordium to the settlement mark on sagittae. This mark was identified as a change in the otolith optical density from darker to opaque (i.e., *Type 1a* settlement mark of Wilson & McCormick, 1999; Fig. 2). Because otolith increments have been validated to be deposited daily in the family Pomacentridae and in some *Stegastes* species (e.g., Wellington & Victor, 1989; Wilson & McCormick, 1999), we assumed that increments were also formed daily in all the species studied here.

**Figure 2.**
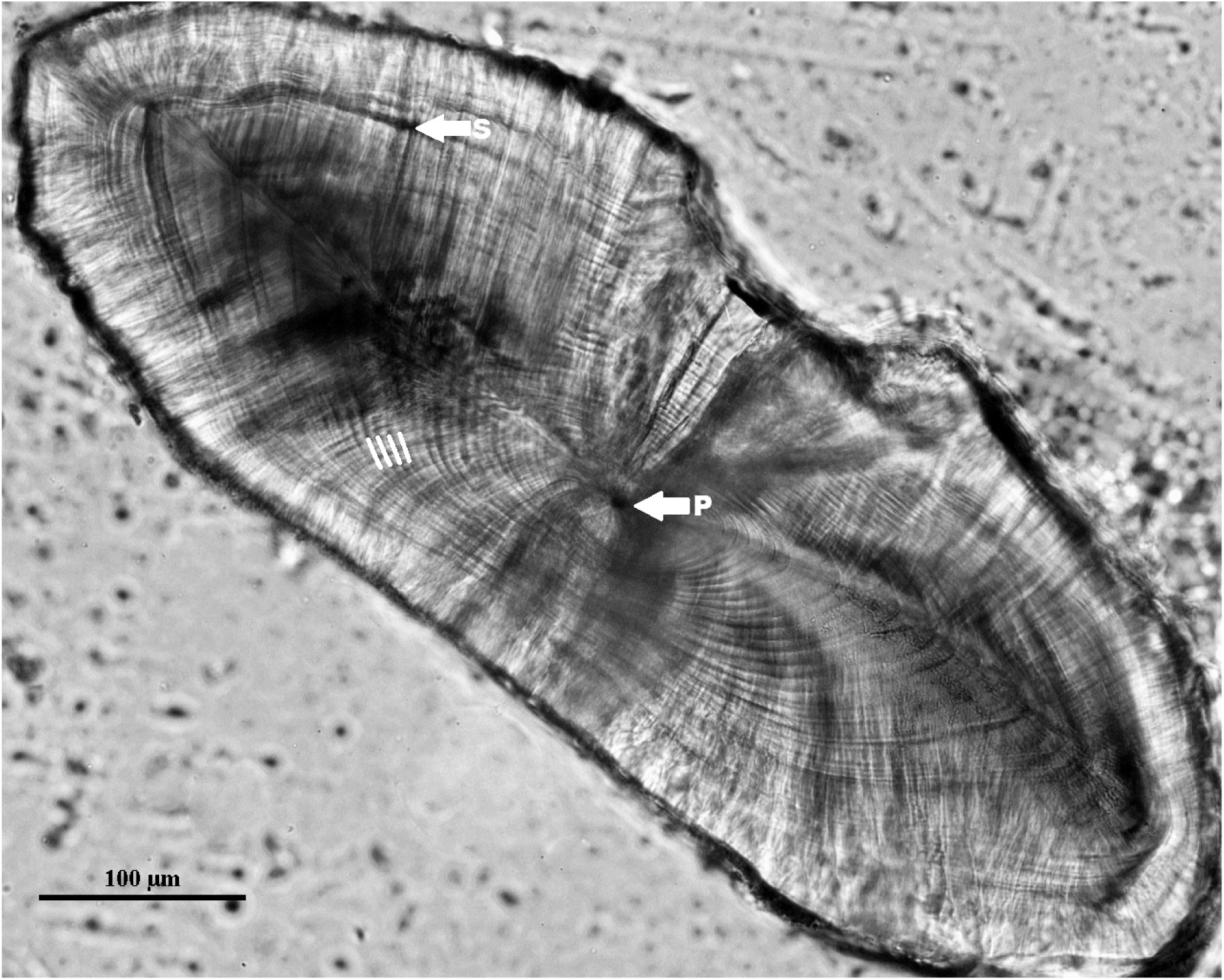
Transverse section of a *Stegastes* sagittae. Solid lines indicate daily increments while arrows indicate the position of the primordium (P) and the settlement mark (S).

The transverse sections of otoliths were examined under an Olympus CH30 microscope and an Eclipse Nikon microscope (40 X magnifications) equipped with a camera and connected to a computer for image analysis (Nis-Basic Research software). Direct readings of the increments under the light microscope were made three times, and only if the counts deviated less than 10% would the mean be used in statistical analyses (Wilson & McCormick, 1999). On the other hand, counts on the microscope with a camera were made using two images of each otolith (one comprising from the primordium to the settlement mark and the other, with higher magnification, around the primordium) to get a better resolution of the increments closer to the primordium. A paired t-test comparing the readings obtained through both microscopes indicated that the images from the computer-equipped microscope showed higher counts, probably due to a better resolution of the increments close to the primordium. To avoid counting sub-daily increments and over-ageing the juvenile fishes, sharpness of otolith projections from the microscope to the computer screen was altered increasing the blurriness to make only the wider/darker increments visible (i.e., daily increments). We used results from the camera-equipped microscope for all the data and analyses reported in this study.

### Statistical Analysis

We examined inter-specific differences in PLD using a one-way ANOVA and planned orthogonal contrasts. Three different sets of mutually orthogonal contrasts were established based in the available degrees of freedom in the “species” factor. One contrast compared the PLD of oceanic species (*S. arcifrons* and *S. beebei*) *vs.* continental species (*S. acapulcoensis* and *S. flavilatus*), another compared the PLD between the two oceanic species (*S. arcifrons vs. S. beebei*) and the last one compared the PLD of the continental species (*S. acapulcoensis vs. S. flavilatus*).

To examine whether inter-specific differences in PLD were consistent between localities, we carried out a two-way ANOVA considering only *S. acapulcoensis* and *S. flavilatus* from three localities (Gorgona Island, Málaga and Utría). Malpelo was not included because only a single *S. flavilatus* was collected there. A Tukey multiple comparison test was used to examine specific differences between different combinations of species and localities. Finally, to examine the consistency of inter-specific differences in PLD at a smaller spatial scale, a two-way ANOVA was also used to compare the same species pair between sites within the same locality (Gorgona Island).

To examine the extent of intra-specific spatial variation in PLD in the continental species we also used one-way ANOVA and planned orthogonal contrasts. For *S. acapulcoensis* (present at all studied localities) three sets of contrasts were performed using the available degrees of freedom in the localities factor. One contrast set compared the PLD estimated from individuals collected at the oceanic island of Malpelo with PLD estimates obtained from the continental localities (Gorgona, Málaga and Utría). Another contrast compared the PLD of individuals from the continental island of Gorgona to that of individuals from the coastal continental localities of Málaga and Utría. The third contrast compared the PLD of individuals collected within coastal continental settings (Utría vs. Málaga). For *S. flavilatus*, we examined differences in PLD among the three continental localities and considered two sets of orthogonal contrasts. The first set compared the PLD of *S. flavilatus* from Gorgona Island with the PLD of individuals from the coastal continental localities (Málaga and Utría). The second contrast compared the PLD of individuals from the two coastal continental localities (Utría vs. Málaga).

In all tests the data complied with assumptions of homoscedasticity and normality, which were examined using Levene’s test (α = 0.05) and inspection of residuals. All statistical analyses were done using Statistica version 8.0 (Statsoft Inc., 2007).

## RESULTS

PLDs varied between 24 and 49 days in the studied damselfish species (Table I). Although there was considerable overlap between their PLD ranges, there were significant differences in mean values among species (ANOVA based on all specimens from all localities, Table IIA). *Stegastes beebei* showed the highest mean PLD followed by *S. arcifrons* and *S. flavilatus*, whereas *S. acapulcoensis* exhibited the lowest (Table I). Orthogonal contrasts revealed statistically significant geographic differences in PLD among the four species. The oceanic species pair (*S. arcifrons* and *S. beebei*) had significantly longer mean PLDs than the continental species pair (*S. acapulcoensis* and *S. flavilatus*) (Table I). Contrasts between the two species within either the continental or the oceanic species pair did not indicate significant differences in mean PLD in either case (Table II).

**TABLE I.**
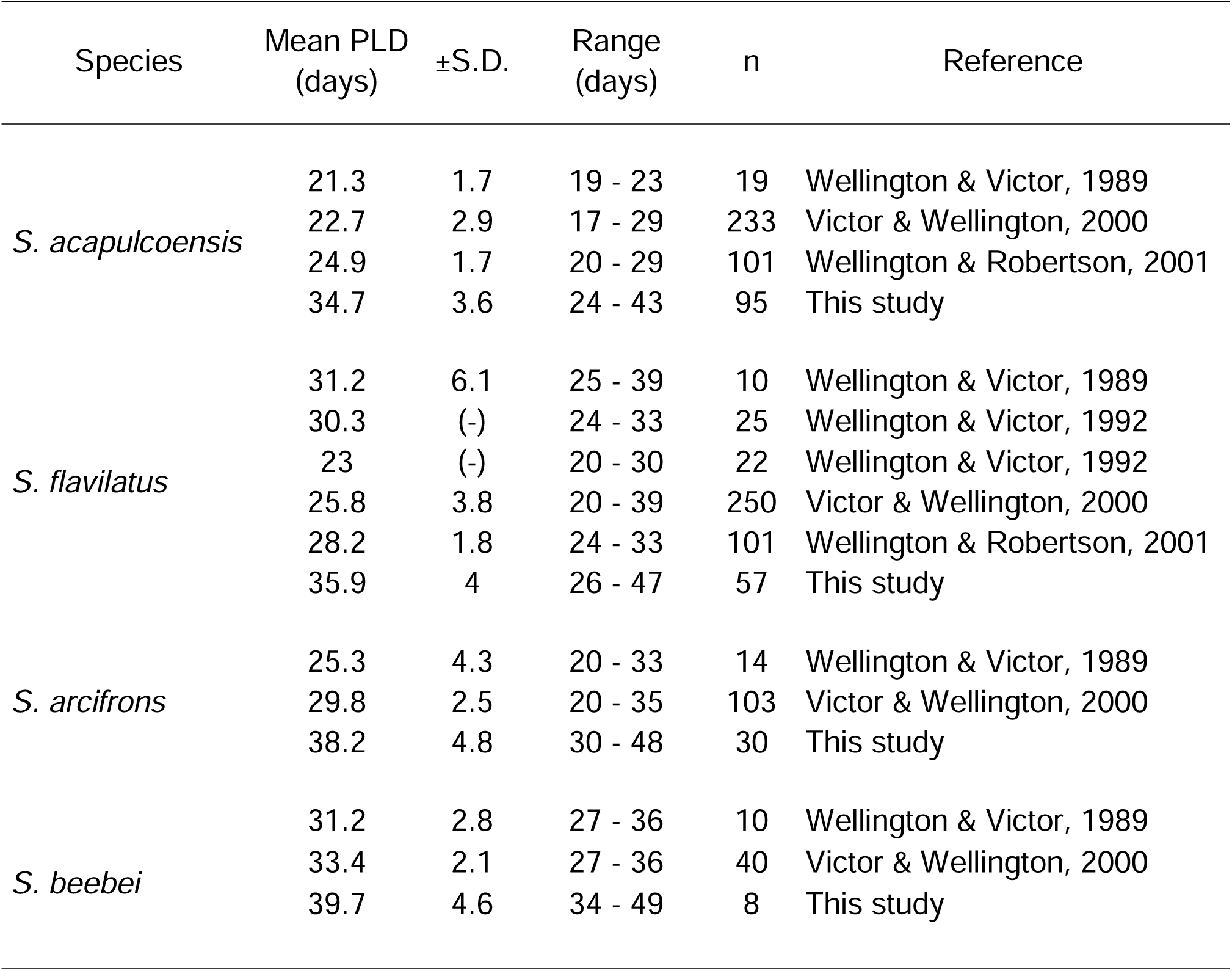
Mean pelagic larval duration (PLD) (±S.D.) in four *Stegastes* species from the Tropical Eastern Pacific. S.D.,Standard deviation; (-), data not available.

**TABLE II.** (A) Results of one-way ANOVA and sets of mutually orthogonal contrasts for inter-specific comparisons (a. oceanic *vs.* continental species; b. among both oceanic species; c. among both continental species). (B) Results of one- way ANOVA testing for intra-specific differences among localities for *Stegastes acapulcoensis* (d. oceanic *vs.* continental localities; e. continental island *vs.* continental coastal localities; f. continental coastal locality *vs.* continental coastal locality). (C) One-way ANOVA testing for intra-specific differences among localities for *S. flavilatus* (g. continental island *vs.* continental coastal localities; h. continental coastal locality *vs.* continental coastal locality). Results of one-way ANOVA and sets of mutually orthogonal contrasts for inter-specific comparisons (a. oceanic *vs.* continental species; b. among both oceanic species; c. among both continental species). Parameter estimates: S.E., Standard error; C.I., Confidence interval.

| (A) Interspecific variation |  | d.f. | SS | MS | *F* | *P* |
| --- | --- | --- | --- | --- | --- | --- |
| Source |  |  |  |  |  |  |
| Species |  | 3 | 405.1 | 135 | 8.63 | 0.000 |
| Error |  | 186 | 2910.9 | 15.6 |  |  |
| Orthogonal contrasts |  |  | SE | t | 95% C.I. | *P* |
| a. (*S. arcifrons*, *S. beebei*) vs. (*S. acapulcoensis*, *S. flavilatus*) |  |  | 1.71 | -4.30 | -10.71 – -3.97 | 0.000 |
| b. *S. arcifrons* vs. *S. beebei* |  |  | 0.66 | -1.72 | -2.45 – 0.17 | 0.087 |
| c. *S. acapulcoensis* vs. *S. flavilatus* |  |  | 1.57 | 0.98 | -1.55 – 4.66 | 0.326 |

Table II (continued)
| (B) Intraspecific variation - *S. acapulcoensis* |  | d.f. | SS | MS | *F* | *P* |
| --- | --- | --- | --- | --- | --- | --- |
| Source |  |  |  |  |  |  |
| Localities |  | 3 | 144.29 | 48.10 | 4.11 | 0.008 |
| Error |  | 91 | 1064.13 | 11.69 |  |  |
| Orthogonal contrasts |  |  | SE | t | 95% C.I. | *P* |
| d. Malpelo vs. (Gorgona, Málaga, Utría) |  |  | 2.72 | 3.02 | 2.82 – 13.64 | 0.003 |
| e. Gorgona vs. (Málaga, Utría) |  |  | 1.71 | 1.58 | -0.69 – 6.09 | 0.117 |
| f. Málaga vs. Utría |  |  | 1.24 | 0.24 | -2.16 – - 2.76 | 0.809 |
| (C) Intraspecific variation - *S. flavilatus* |  | d.f. | SS | MS | *F* | *P* |
| Source |  |  |  |  |  |  |
| Localities |  | 2 | 228.08 | 114.04 | 9.84 | 0.000 |
| Error |  | 53 | 614.42 | 11.59 |  |  |
| Orthogonal contrasts |  |  | SE | t | 95% C.I. | *P* |
| g. Gorgona vs. (Málaga, Utría) |  |  | 2.09 | -4.32 | -13.24 – -4.85 | 0.000 |
| h. Málaga vs. Utría |  |  | 1.76 | 0.91 | -1.93 – 5.13 | 0.367 |

The magnitude of differences in PLD between *S. acapulcoensis* and *S. flavilatus* was not constant at all localities (Fig. 3). In particular, mean PLD of these two species was similar at Gorgona Island, but significantly different at the other two coastal localities. The PLD of the single individual of *S. flavilatus* collected at Malpelo Island was relatively long (43 days; Table III), but Malpelo was not included in the statistical comparisons for lack of replication in this species.

**Figure 3.**
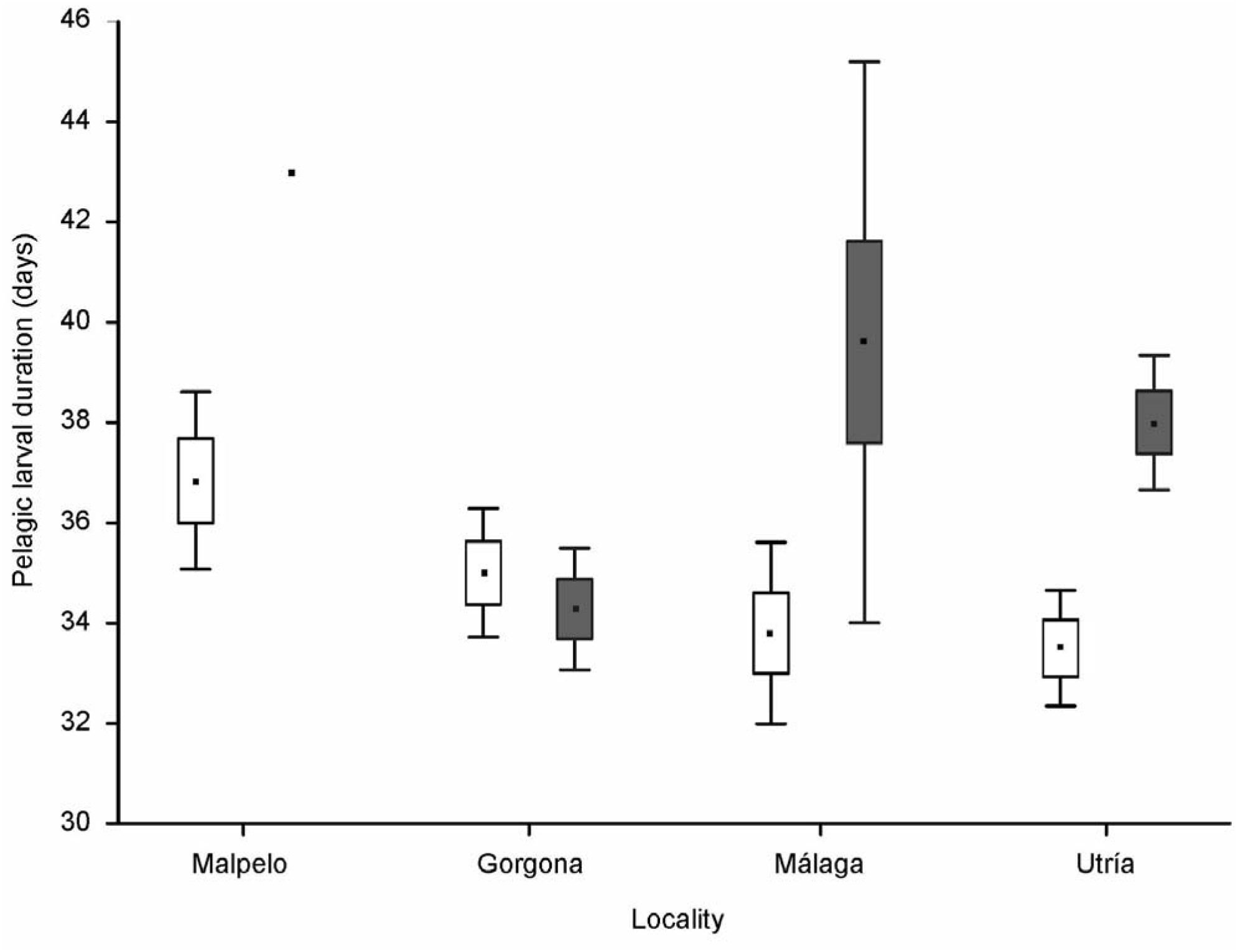
Spatial variation in the pelagic larval duration of *Stegastes acapulcoensis* (white boxes) and *S. flavilatus* (grey boxes) at different localities (oceanic island, continental island, mainland locations) in the Colombian Pacific. Whiskers represent 95% confidence intervals and boxes one SE from the mean.

**TABLE III.** Mean pelagic larval duration (PLD) (±S.D.) for four species of damselfish (genus *Stegastes*) at an oceanic island (Malpelo), a continental island (Gorgona) and two continental coastal localities (Málaga and Utría) from the Colombian Pacific. Estimates were based on the increments counts in the sagittae. S.D., Standard deviation; (-), Standard deviation could not be calculated.

| Species | Location | n | Mean PLD<br>(days) | ±S.D. | Range<br>(days) |
| --- | --- | --- | --- | --- | --- |
| *S. acapulcoensis* | Malpelo | 19 | 36.8 | 3.67 | 30 – 43 |
|  | Gorgona | 34 | 35 | 3.67 | 24 – 42 |
|  | Málaga | 10 | 33.,8 | 2.53 | 28 – 37 |
|  | Utría | 32 | 33.5 | 3.2 | 28 – 41 |
| *S. flavilatus* | Malpelo | 1 | 43 | (-) | 43 |
|  | Gorgona | 36 | 34.2 | 3.59 | 26 - 42 |
|  | Málaga | 5 | 39.6 | 4.5 | 36 - 47 |
|  | Utría | 15 | 38 | 2.42 | 34 - 43 |
| *S. arcifrons* | Malpelo | 30 | 38.2 | 4.76 | 30 - 48 |
| *S. beebei* | Malpelo | 8 | 39.8 | 4.62 | 34 - 49 |

There was significant intra-specific spatial variation in PLD in both *S. acapulcoensis* and *S. flavilatus* - the latter being the only species collected at more than one locality (Tables II and III; Fig. 3). For *S. acapulcoensis*, the larval duration of individuals from the oceanic island of Malpelo was significantly longer than that of individuals from continental localities (Gorgona, Málaga and Utría; Table IIB).

However, individuals from the continental island (Gorgona) did not differ from those of coastal continental localities (Málaga and Utría), and individuals from the latter two localities did not differ from each other. Nonetheless, *S. acapulcoensis* showed a trend of decreasing PLD from the offshore oceanic locality towards the mainland coastal localities (Table III). In the case of *S. flavilatus*, individuals from Gorgona Island had a significantly shorter PLD than those from the coastal localities, but individuals from the coastal localities did not differ in PLD (Tables IIC and III).

Finally, no significant intra-specific differences in PLD were found between sites within a locality. The mean PLD of *S. acapulcoensis* did not differ from that of *S. flavilatus* either at the coral reef of La Ventana or the rocky reef of El Laberinto at Gorgona Island (Fig. 4).

**Figure 4.**
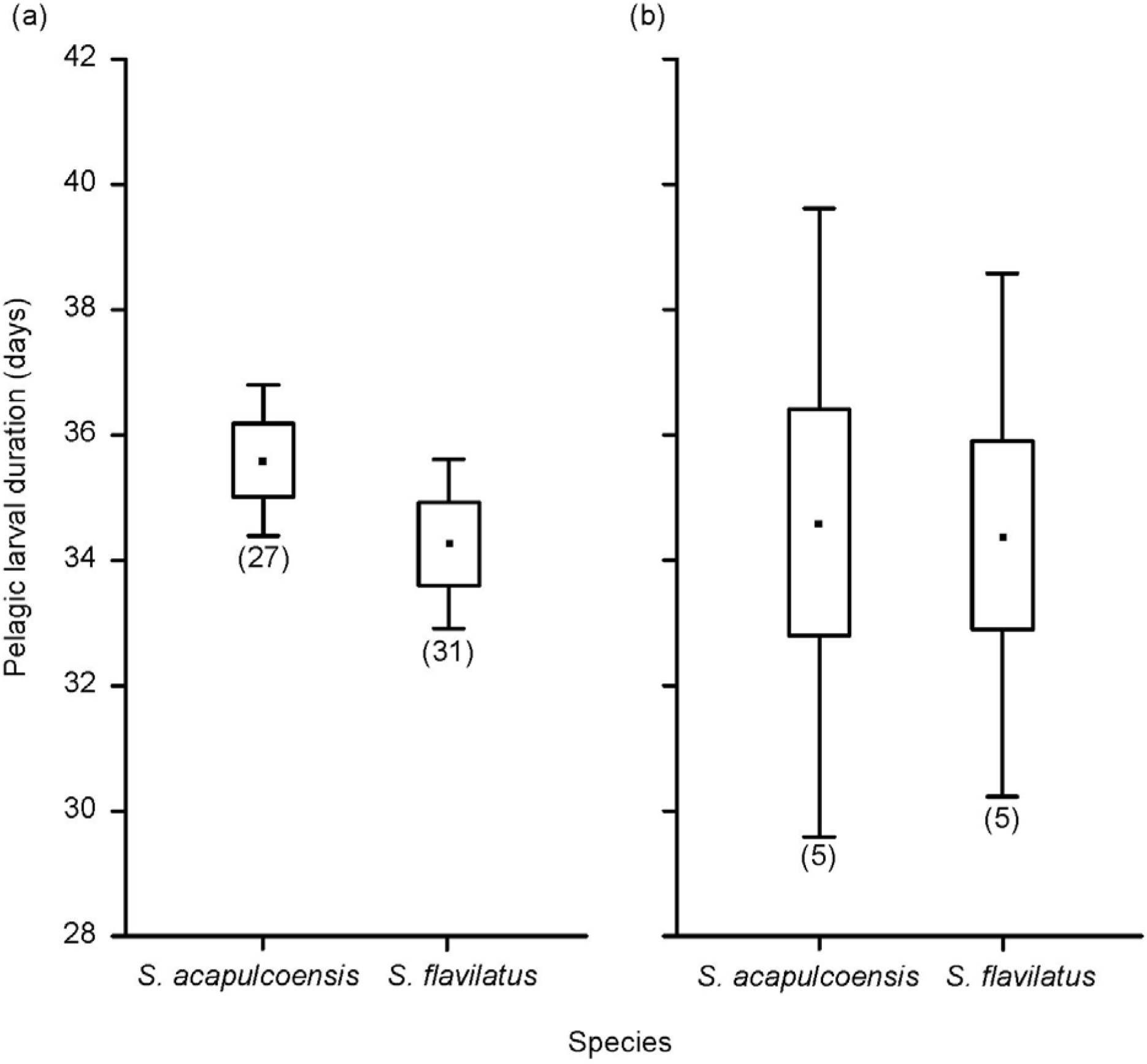
Pelagic larval duration for *Stegastes acapulcoensis* and *S. flavilatus* at two sites within Gorgona Island: (a) El Laberinto and (b) La Ventana. Whiskers represent 95% confidence intervals and boxes one SE from the mean.

## DISCUSSION

This study provides evidence of significant inter- and intra-specific variation in the PLD of four damselfish species (*Stegastes* spp.) from the TEP between localities within the Colombian Pacific (scale of 100’s km), but not between sites within a locality (10’s km). Much of the variation in PLD observed in this study was related to the oceanic *vs.* continental distribution of species or individuals.

While differences in PLD between congeneric species are not unusual in reef fishes in general and damselfishes in particular (Thresher *et al*., 1989; Wellington & Victor, 1989; Victor & Wellington, 2000; Robitzch et al. 2016), in this study we document a seemingly predictable pattern: species with an oceanic distribution as a group (*S. arcifrons* and *S. beebei*) showed longer larval durations than the species with primarily continental geographic ranges (*S. acapulcoensis* and *S. flavilatus*). This pattern is reinforced by the fact that mean PLD did not differ between these two species pairs. Victor & Wellington (2000) had previously found similar results for damselfishes in the TEP, but in different localities than those considered in the present study. They reported longer PLDs for the endemic insular species at Cocos, Clipperton and the Galapagos Archipelago compared to the PLDs of their widely distributed congeners sampled along the Pacific coast of Baja California and Panama. When considering the PLD for all damselfish species in the TEP it seems like this is a general pattern except for *S. rectifraenum* and *S. baldwini* (Table I). It has been noted that fish species at isolated localities can have significantly longer PLDs than congeneric species at larger and well-connected localities (Brother & Thresher, 1985; Victor, 1986; Cowen & Sponaugle, 1997).

Contrary to expectations regarding an enhanced probability of larval retention by a short PLD in remote localities (Robertson, 2001), it would seem that a longer rather than a shorter PLD is associated with population maintenance at isolated oceanic localities. Evidence supporting this hypothesis has been found in endemic reef fishes on isolated islands (Hawaiian Islands in the Pacific Ocean and Christmas and Cocos-Keeling Islands in the Indian Ocean) whose larvae have longer PLDs and adults are found in high abundances (Victor, 1986; Hourigan & Reese, 1987; Hobbs, 2011; Hobbs *et al*., 2011).

For insular isolated species, longer PLDs may be a requirement to increase the probability of finding suitable habitat. Indeed, for endemic species longer PLDs increase the probability of reaching their natal reef after navigating current systems. A longer larval duration possibly gives them the opportunity to reach other oceanic islands, which would imply the existence of high connectivity between the oceanic islands in the southern TEP (Malpelo, Cocos, Galápagos). However, little is known about the connectivity in marine organisms between islands in this region. Another alternative is the occurrence of self-recruitment with a long pre-settlement period. This scenario might be occurring in Malpelo, which has also been demonstrated for damselfishes in other regions (Cowen & Castro, 1994; Kingsford *et al*., 2011). The oceanographic and physical characteristics of coral reefs on oceanic islands (e.g., narrow reef platform, energetic currents) may favor a rapid and large dispersal from their natal reef (Cowen, 2002). Homing to natal reefs has been explained as a result of local circulation patterns and vertical migration of larvae, which allows them to use currents in different directions and avoid predation (Lecchini et al., 2013). Evidence of this scenario was reported for damselfishes in Barbados and Papua New Guinea (Cowen & Castro, 1994; Almany, 2007).

The level of isolation and endemism in Malpelo suggests that the larvae of several groups of organisms are returning to their natal reefs. In the case of *Stegastes* spp., it is possible that a longer PLD increase the probability of larvae returning to the island. In a remote environment the chances of finding a suitable location to settle is diminished significantly if larvae are dispersed beyond their parental population (Swearer *et al*., 2002). Studies on the genetic population structure of these fish populations could therefore help to provide insights on the dispersal dynamics in the marine populations of this region (Rodríguez-Moreno et al. 2017). In order to understand the occurrence of self-recruitment or high connectivity in damselfish populations it is important to carry out studies that consider the natal reef of the larvae, using techniques such as otolith chemistry and genetic markers, to allow an accurate discrimination of larval sources (as in Thorrold, 2006; Saenz-Agudelo *et al*., 2011; Berumen *et al*., 2012, Harrison *et al*., 2012).

The intra-specific variations in PLDs among localities observed in *S. flavilatus* and *S. acapulcoensis* agrees with other studies based in the TEP and on the Great Barrier Reef (Wellington & Victor, 1992; Bay *et al*., 2006), where spatial variability in the PLD of Pomacentrid fishes has also been reported. In our study, spatial variation of PLD revealed that the longer PLD estimates in *S. acapulcoensis* was from individuals collected at the isolated island of Malpelo. The existence of long-range dispersal events for fertilized eggs that in distant locations may explain this observation. Other mechanisms that could help enhance fish dispersal are the delay of metamorphosis and the transport by rafting with floating objects (Victor, 1986; Kokita & Omori, 1999; Mora *et al*., 2001). The PLD of *S. acapulcoensis* in Malpelo ranged from 30 to 43 days. Based on the apparent absence of a reproductively viable population of this species on Malpelo (MRM pers. observations), the most probable source of these larvae is from the continent, which is located 365 km away from the island. Victor (1987) reported a range of PLDs (23 to 38 days) for some *Stegastes* spp. in the eastern Pacific that were found several hundreds of kilometers offshore. This observed dispersal potential may not be representative of damselfishes in general, but could be of great importance for rare colonization events recorded in the TEP.

The colonization of some islands by *S. acapulcoensis* is striking. Before 1982 this damselfish was considered a species restricted to continental areas (Grove *et al*., 1986). After the El Niño of 1982-1983, individuals of this species were recorded in the Galapagos Islands (Grove *et al*., 1986; Victor *et al*., 2001). According to Robertson & Allen (2015), its range now includes Galapagos, Revillagigedos, Cocos and Malpelo islands. The presence of some adults of *S. acapulcoensis* and the frequent arrival of juveniles to Malpelo indicates the potential for colonization of this species. *Stegastes flavilatus*, however, has not been that successful colonizing islands, although there are records of a few individuals in some islands (Grove & Lavenberg, 1997; Rodríguez-Moreno, 2011). This observation is intriguing given the greater mean and range in the PLD of *S. flavilatus* in comparison to *S. acapulcoensis*.

Our PLD estimates for all the species studied here were higher than previous reports. It is possible that fishes in the southern portion of the TEP naturally exhibit longer PLD’s given that none have yet been reported in the Colombian Pacific.

Evidence supporting this hypothesis can be found in records of *S. flavilatus*, which had a PLD of 27 days in Panama (central portion of the TEP) and increased to 35 days in individuals collected farther south in Ecuador (Victor & Wellington, 1989). Also Meekan *et al*. (2001) reported differences in their demography between populations in the center versus the edges of their geographic distribution.

However, these higher values of PLD should be further examined.

Recent studies reported a higher variation of PLD among sites that were geographically closer to each other than among sites further apart (Di Franco & Guidetti, 2011; Kingsford *et al*., 2011). In our study, we did not find differences in PLD among sites for either *S. acapulcoensis* or *S. flavilatus* at Gorgona Island.

Small spatial scale variability in PLD has been associated with environmental conditions and small-scale oceanography (Di Franco & Guidetti, 2011). Because oceanographic conditions at Gorgona Island vary within a year, with two contrasting periods (May-December: warm and low salinity; January-April: cold and high salinity: Giraldo *et al*., 2008), our results cannot be generalized as we did not consider this seasonal component.

In conclusion, this study confirms variation in the PLD for species of *Stegastes* in the Colombian Pacific. Our results help to understand the dispersal potential of damselfishes (using PLD as a proxy for dispersal ability), which is a key piece of information to the development of conservation strategies. Although individuals of continental species exhibit PLDs that would allow them to reach oceanic habitats and vice-versa, we did not observe any successful colonization events, suggesting that their distribution is not limited by dispersal but other processes instead. For instance, phylogeography studies on some *Acanthurus* species in the Atlantic near to the Amazon River plume show that when oceanic distances are not an obstacle to larval dispersal their distribution was driven by adult specific habitat preferences (Rocha *et al*., 2002). Robitzch et al. (2016) found that PLDs for the endemic damselfishes in the Red Sea (*Dascyllus*) were significantly correlated with food availability, latitude, and temperature. Also a recent study states that adult life-history traits (body size, schooling behavior, and nocturnal activity) affecting recruitment and post-settlement survival might be key determinants of geographic range size in reef fishes (Luiz *et al*., 2013). However, population maintenance and gene flow (i.e., local replenishment or long-distance dispersal) of *Stegastes* in the Colombian Pacific and in the TEP is in need of further examination.

The authors want to express their gratitude to C. Mora (University of Hawaii) and P. Herrón for instructing D. F. Lozano-Cortés and M. Rodríguez-Moreno in otolith extraction and processing, respectively. C. Muñoz, M. López-Victoria, E. Florez and J. Daza helped during field collections and P. Mejía-Falla and A. Navia from Fundación SQUALUS provided advice on image processing. The draft of this manuscript was improved thanks to comments by M.M. Palacios, M. López-Victoria, H.B. Harrison, P. Saenz-Agudelo, M. Berumen, J. DiBattista and the members of the Reef Ecology Lab at King Abdullah University of Science and Technology. Also we want to thank UAESPNN (Colombian National Parks), the Coral Reef Ecology Research Group, the Science Postgraduate Image Laboratory of Universidad del Valle and Juan Felipe Ortega. Many thanks to Fundación Malpelo y Otros Ecosistemas Marinos for logistical support. Financial support for this study was provided by Universidad del Valle and the Administrative Department of Science, Technology and Innovation of Colombia (COLCIENCIAS) through the program “Jóvenes Investigadores e Innovadores - Virginia Gutiérrez de Pineda 2011” awarded to D. Lozano-Cortés, and a scholarship for doctoral studies awarded to M. Rodríguez-Moreno.

## Notes

### Competing Interest Statement

The authors have declared no competing interest.

